# Obesity and adiposity have opposing genetic impacts on human blood traits

**DOI:** 10.1101/2021.11.05.467482

**Authors:** Christopher S Thom, Madison B Wilken, Stella T Chou, Benjamin F Voight

**Author notes:** Correspondence: Christopher S Thom, Children’s Hospital of Philadelphia, Abramson Research Center, Room 415, 3615 Civic Center Blvd, Philadelphia, PA 19104, Benjamin F Voight, Smilow Center for Translational Research, Room 10-126, 3400 Civic Center Blvd, Philadelphia, PA 19104.

## Abstract

Obesity, hyperlipidemia, and truncal adiposity concordantly elevate cardiovascular disease risks, but have unknown genetic effects on blood trait variation. Using Mendelian randomization, we define unexpectedly opposing roles for generalized obesity and truncal adiposity on blood traits. Elevated genetically determined body mass index (BMI) and lipid levels decreased hemoglobin and hematocrit levels, explaining consistent with clinical observations associating obesity and anemia. However, lipid-related effects were confined to erythroid traits, whereas BMI affected multiple blood lineages, indicating broad effects on hematopoiesis. BMI-related effects were unexpectedly opposed by truncal adipose distribution, which increased hemoglobin and blood cell counts across lineages. Conditional analyses indicated genes, pathways, and cell types responsible for these effects, including *Leptin Receptor* and other blood cell-extrinsic factors in adipocytes and endothelium, which regulate hematopoietic stem and progenitor cell biology. Our findings identify novel roles for obesity and adipose distribution on hematopoiesis and show that genetically determined adiposity plays a previously underappreciated role in determining blood cell formation and function.

## Introduction

Blood cell homeostasis is achieved through incompletely understood coordination of blood cell-intrinsic gene regulation and blood cell-extrinsic environmental mechanisms (Comazzetto et al., 2021; Ulirsch et al., 2019). The importance of blood cell formation and function in normal blood development, hematologic disease pathology, and clinical manifestations of systemic disorders has prompted extensive investigation of loci underlying human blood trait variation through genome wide association studies (GWAS) (Astle et al., 2016; Chen et al., 2020; Vuckovic et al., 2020). However, it has been difficult to identify extrinsic effects within these data.

Adipocytes and endothelial cells within the bone marrow environment regulate hematopoiesis (Comazzetto et al., 2021; Zhong et al., 2020). However, discrete adipocyte populations differentially modulate systemic physiology and homeostasis (Hildreth et al., 2021). For example, white adipose tissue has a derogatory effect on hematopoiesis, whereas mesenchymal-derived bone marrow adipocyte populations support blood cell formation (Comazzetto et al., 2021; Cuminetti, 2019; Wang et al., 2018; Zhong et al., 2020).

Genetic studies have revealed that obesity increased metabolic and cardiovascular disease risks (Pulit et al., 2019), and that adipose distribution influences these effects (Huang et al., 2021). While some observational studies have linked obesity (Aigner et al., 2014) or hypercholesterolemia (Shalev et al., 2007) with anemia, others observed apparent erythrocytosis in obese patients (Keohane et al., 2013). Genetic relationships have not been elucidated for adiposity (waist-to-hip ratio, WHR) or obesity (body mass index, BMI) on erythroid or other blood traits.

Mendelian randomization (MR) leverages variants linked to an exposure trait to estimate causal genetic effects on an outcome (Hemani et al., 2018). Multivariable MR (MVMR) and causal mediation analyses can parse effects from multiple factors (Burgess et al., 2017). Using an MR framework, we interrogated causal effects of obesity and obesity-related factors on erythroid and other blood traits, revealing unexpected associations that point to unexpected biological factors and pathways between these clinically relevant human traits. Conditional genome-wide analyses using mtCOJO (Zhu et al., 2018) highlighted loci that were substantially influenced by obesity and adiposity distribution, helping to reveal genes and pathways by which these physiology factors impact blood trait variation.

## Results

Using MR, we found that each standard deviation (SD) unit increase in BMI caused a 0.057 SD decrease in hemoglobin (HGB) levels by the inverse variance weighted method (p=1.0×10^−5^) that was directionally consistent across sensitivity analyses without evidence of horizontal pleiotropy or weak instrument bias (**Figure 1a, Figure 1-figure supplement 1a, Supplementary file 1-Table 1**). Similar effects were observed for hematocrit (HCT, **Figure 1b** and **Figure 1-figure supplement 1b**), suggesting BMI is genetically linked with a lower hemoglobin.

**Figure 1.**
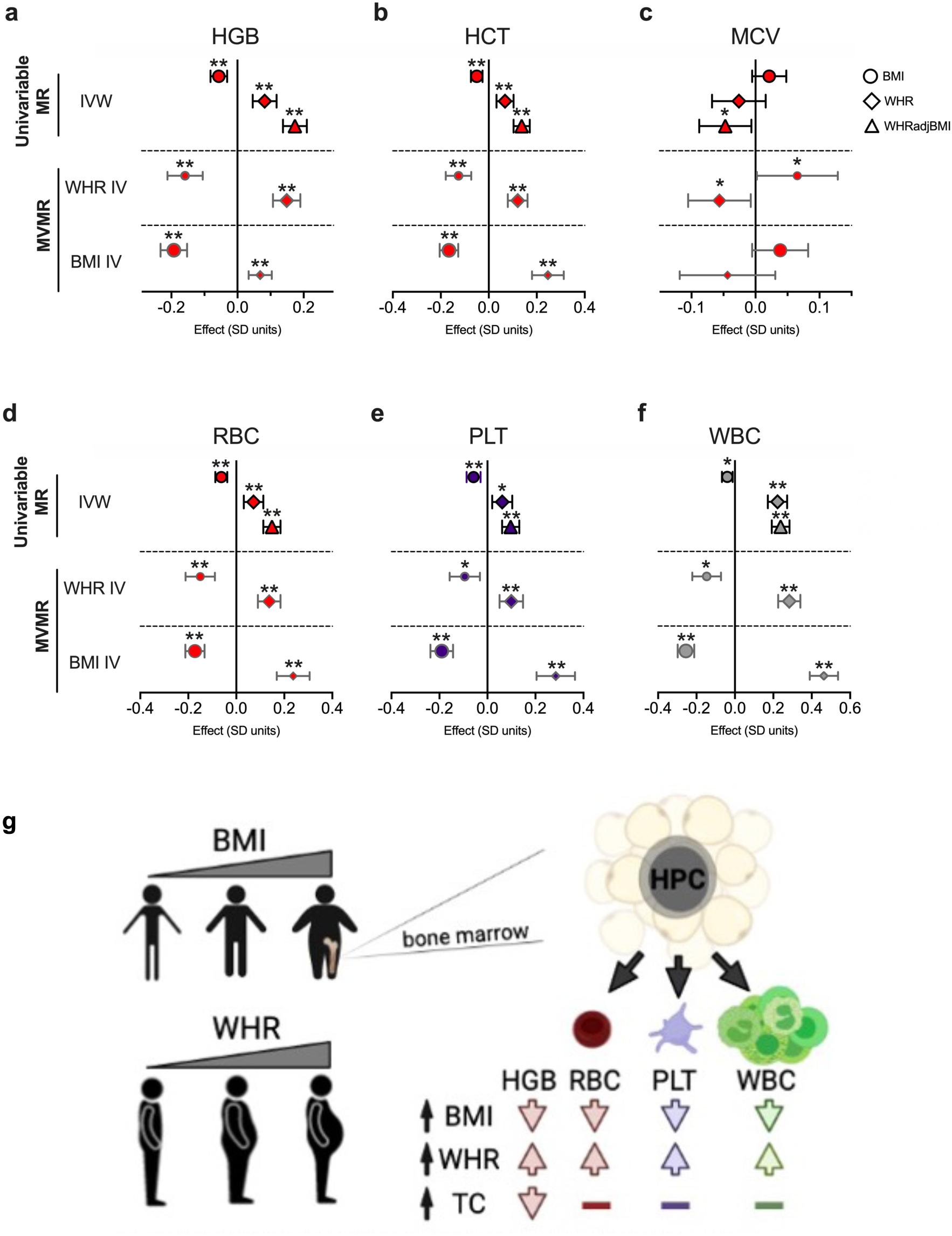
BMI and WHR exert opposing effects on erythroid traits. **(a-f)** Effects of BMI, WHR, WHRadjBMI on (**a**) hemoglobin, HGB, (**b**) hematocrit, HCT, or (**c**) mean corpuscular volume, MCV, (**d**) red blood cell count, RBC, (**e**) platelet count, PLT, or (**f**) white blood cell count, WBC. Shown in top panel are effects of BMI, WHR, or WHRadjBMI on HGB in univariable MR experiments by inverse variance weighted method (IVW). Effects are in SD units with 95% confidence intervals. Underneath univariable MR results, effects of BMI or WHR at 639 LD-independent WHR-associated SNPs are shown. Bottom row of panels show effects of BMI or WHR at 1268 LD-independent BMI-associated SNPs. *p<0.05, **p<0.003. (**g**) Schematic summarizing effects of indicated exposures on blood traits.

We next investigated previously proposed mechanisms to explain observational links between BMI and anemia. For example, hypercholesterolemia may cause anemia (Shalev et al., 2007) through erythrocyte membrane formation and stability (Mohandas and Gallagher, 2008). We confirmed that increased total cholesterol (TC) or lipid fractions (low density lipoprotein or high density lipoprotein), but not triglyceride levels, decreased hemoglobin or hematocrit (**Figure 1- figure supplement 2-3**). However, multivariable mediation experiments revealed that cholesterol levels alter erythroid traits via mechanisms independent from BMI (**Figure 1-figure supplement 3b** and **Supplementary file 1-Table 2**). Chronic inflammation and iron deficiency, which cause decreased erythrocyte size (microcytosis), have also been suggested to mediate obesity-related anemia (Aigner et al., 2014). However, BMI did not alter erythrocyte mean corpuscular volume (MCV) by MR (p=0.12, **Figure 1c, Figure 1-figure supplement 4a**). These findings aligned with clinical observations linking BMI with anemia risk, but argued against prevailing mechanistic hypotheses at the genetic level.

Reverse causality experiments also identified inverse correlations between erythroid and metabolic traits (**Figure 1-figure supplement 5**). Directional MR Steiger (Hemani et al., 2017) analyses were inconsistent (**Supplementary file 1-Table 3**), perhaps limited by blood trait measurement variation or quantitative adjustments for individual characteristics (Vuckovic et al., 2020).

We then considered an alternative hypothesis that the physiological distribution of adipose as measured by WHR could impact obesity-related anemia risk. Unexpectedly, and in contrast to BMI, higher WHR increased red blood cell traits (**Figure 1a,b** and **Figure 1-figure supplement 6a,b**), with WHR adjusted for BMI on an individual level (WHRadjBMI) exacerbating these positive effects (**Figure 1a,b** and **Figure 1-figure supplement 6c,d**) while also associating with decreased MCV (**Figure 1c** and **Figure 1-figure supplement 4b,c**). Multivariable analyses formally validated the opposing, cross-mediating effects of both measures of adiposity on erythroid traits (**Figure 1a,b,c**).

Multivariable and mediation analyses on other blood traits identified cross-mediating opposing effects of WHR and BMI on quantitative blood counts across cell lineages (**Figure 1d,e,f** and **Figure 1-figure supplement 7-8**). These effects persisted after accounting for related blood traits (**Figure 1-figure supplement 9**). The directionally consistent effects across multiple lineages suggested that underlying mechanisms related to hematopoietic stem and progenitor cells (HSCs) common to these lineages (Thom and Voight, 2020) (**Figure 1g**). These findings also argued against gender-related effects: although men generally have higher hemoglobin (Vuckovic et al., 2020) and WHR (Pulit et al., 2019), women have higher platelet and neutrophil counts (Bain, 1996).

Next, we used mtCOJO applied to blood trait GWAS data to condition on polygenic-measured BMI and/or WHR to identify blood loci related to these factors (Zhu et al., 2018). BMI adjustment modestly changed SNP effect sizes and retained most (91%) unadjusted lead sentinel associated variants (**Figure 2a,b** and **Supplementary file 1-Table 4**). However, combined BMI and WHR adjustment resulted in substantial SNP effect size changes (>36-fold increased standard deviation vs BMI adjustment alone, p<0.0001 by F test to compare variances), with a negative skew (−2.0) reflecting adjustment for the positive effect of WHR on hemoglobin levels (**Figure 2a,b**). Thus, while prior GWAS adjusted for BMI (Astle et al., 2016), we identified more prominent effects for WHR across blood traits and lineages.

**Figure 2.**
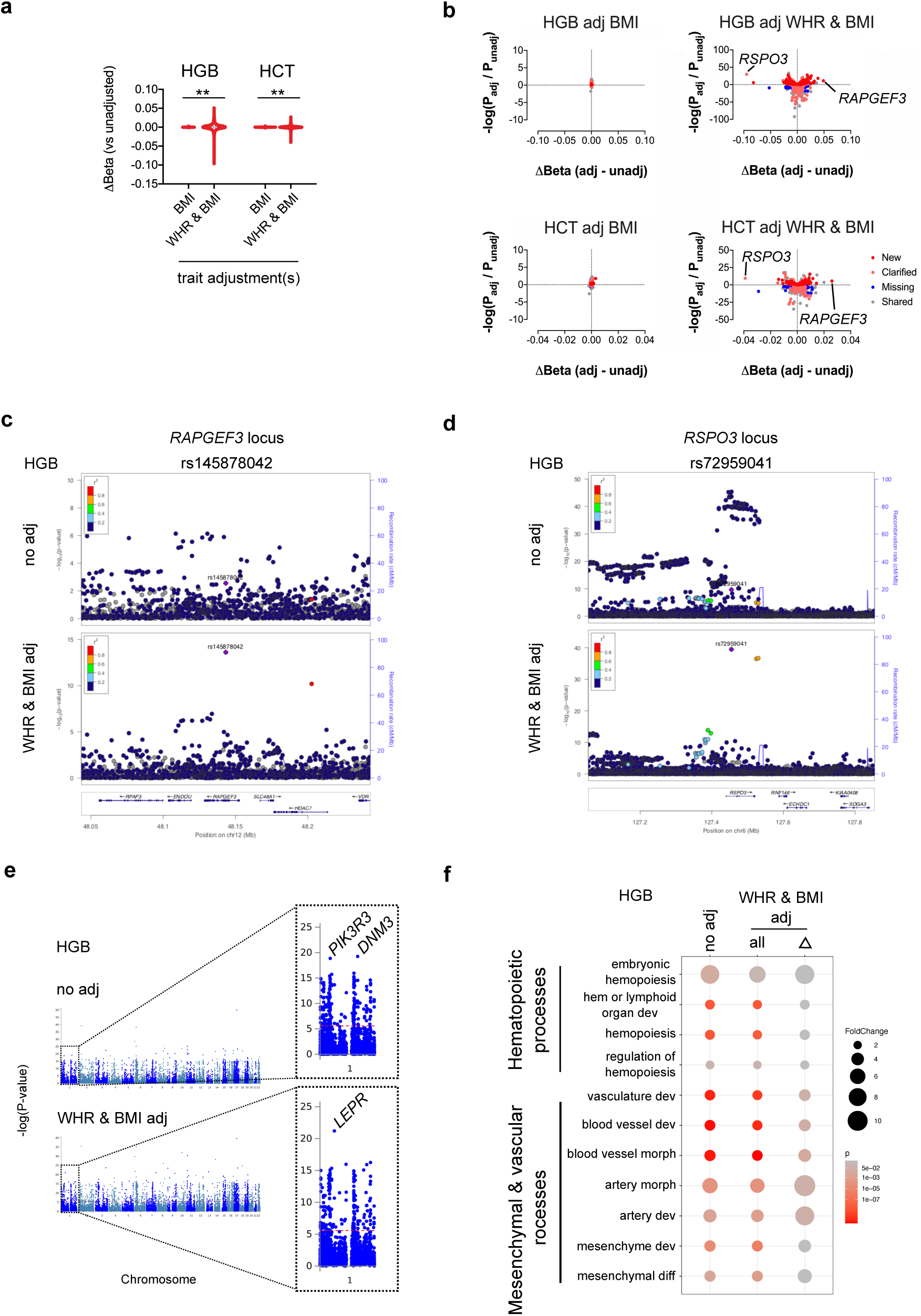
Conditional blood trait analysis based on obesity and/or adiposity modifies interpretation of genomic loci that impact blood trait variation. (**a**) Violin plots showing the dispersion in effect size at genome-wide significant loci after adjusting erythroid traits (HGB or HCT) for BMI, or WHR and BMI. **p<0.0001 by F test to compare variances. (**b**) Scatterplots depicting changes in effect sizes and p-values for all genome-wide significant sentinel loci before or after adjustment. Novel loci (red) had p<5×10^−8^ only after adjustment and represent new loci (not in LD with genome-wide significant SNPs before adjustment). Clarified loci (pink) are sentinel SNPs with p<5×10^−8^ after adjustment and are in linkage disequilibrium with significant pre-adjustment SNPs. Missing loci (blue) are those with adjusted p>5×10-8, which were significant pre-adjustment. Shared SNPs (gray) are sentinel SNPs before and after adjustment for the indicated factors. (**c**) After adjustment for WHR and BMI, the common coding SNP (rs145878042) in *RAPGEF3* significantly impacts HGB level. (**d**) Adjustment for WHR and BMI alters interpretation of SNP effects at the *RSPO3* locus, including more significant effects for new sentinel variant rs72959041 (unadjusted p=2.1×10^−10^, adjusted p=3.4×10^−40^). (**e**) Gene-based Manhattan plots for HGB, before or after BMI/WHR adjustment. (**f**) Gene ontology analyses for hematopoietic, mesenchymal, and vascular biological processes for HGB loci before and after mtCOJO adjustment for BMI and WHR. Significance reflects Fisher’s exact test after multiple testing.

Combined BMI/WHR adjustment shifted 341 hemoglobin-associated loci toward the null, supporting a key role for obesity- or adiposity-mediated mechanisms at these sites while also identifying 844 hemoglobin loci that either clarified interpretation of previously implicated loci or tagged previously unreported regions (n=242 loci, **Figure 2b** and **Supplementary file 1-Table 5**). For example, a missense coding variant in *RAPGEF3* (rs145878042), previously linked to obesity (Pulit et al., 2019), adiposity (Pulit et al., 2019), and platelet distribution width (Vuckovic et al., 2020), did not meet genome-wide significance for hemoglobin (p=0.003) until BMI/WHR adjustment (p=2.4×10^−14^, **Figure 2b,c**). Interpretation of SNPs at the *RSPO3* locus also dramatically changed (**Figure 2b,d**). *RSPO3*, a Wnt pathway modulator that directs development of bone and other tissues (Nilsson et al., 2021), has been linked with adiposity (Pulit et al., 2019) and blood trait variation (Vuckovic et al., 2020). Similar effects were seen in adjusted hematocrit data (**Figure 2a,b** and **Supplementary file 1-Tables 6-7**) and quantitative traits across blood lineages (**Figure 2-figure supplement 1**). These conditional analyses presumably revealed sites where BMI and/or WHR biology most strongly impact blood trait variation, although it is possible that some pleiotropic loci independently regulate blood traits through shared or different gene regulation.

Functional enrichment analyses identified many consistent genes and processes in unadjusted versus BMI/WHR-adjusted data across blood traits, with some notable changes (**Figure 2e,f Figure 2-figure supplement 2** and **Supplementary file 1-Tables 8-23**) (Mi et al., 2019; Watanabe et al., 2019). For example, at the gene level, association at *LEPR* in these adjusted analysis for HGB elevated to statistical attention (**Figure 2e**). *LepR* expression impacts obesity predisposition, and LepR^+^ endothelial niches support HSC survival (Comazzetto et al., 2021). Further, adjusted HGB locus-related genes were enriched for some endothelial and mesenchymal development processes, albeit with diminished p-values from power loss from limited SNP sets (**Figure 2f** and **Supplementary file 1-Tables 8-17**). Adjusted RBC, PLT, and WBC data also demonstrated enrichment of endothelial and cell adhesion pathways (**Figure 2- figure supplement 1** and **Supplementary file 1-Tables 18-23**). These findings highlight the relevance for adiposity and related biology in regulating multilineage blood traits, including contributions from mesenchyme-derived adipocytes (Zhong et al., 2020) and stromal endothelial cells in bone marrow (Comazzetto et al., 2021). Indeed, adipose tissue-related genes were enriched in multilineage blood trait data (**Figure 2-figure supplement 2**).

## Discussion

The obesity epidemic has increased the importance of understanding associated systemic comorbidities (Koenen et al., 2021), including complex physiology linking cardiometabolic and blood traits (Comazzetto et al., 2021). While some clinical epidemiological studies have proposed iron deficiency and chronic inflammation to explain the anemia observed in some obese populations (Aigner et al., 2014; Benova and Tencerova, 2020; Koenen et al., 2021), confounders inherent to observational studies may limit interpretation. Our results show that genetically determined BMI is indeed causally associated with lower hemoglobin and hematocrit levels. However, we identified multilineage regulation that extended beyond clinically reported obesity-related effects, which may facilitate laboratory result interpretation and clinical management in the right context.

Further, our studies suggest that obesity and adiposity act through different mechanisms than previously proposed (Aigner et al., 2014; Koenen et al., 2021). Directionally consistent effects across blood lineages may indicate obesity- and adiposity-related influences on hematopoietic stem and progenitor cells (HSCs) in the bone marrow. A genetic predisposition to accumulate bone marrow fat in the form of white adipose tissue may underlie age- (Tuljapurkar et al., 2011) or obesity-related (Benova and Tencerova, 2020) cytopenias by regulating HSC self-renewal or differentiation (Wang et al., 2018). Genetically determined differences in bone marrow stromal cell types (e.g., mesenchymal stem cell-derived bone marrow adipocytes (Zhong et al., 2020)) could also impact HSC biology. It is possible that decreased hip circumference, manifested as *increased* WHR, could reflect decreased predisposition to accumulate HSC-inhibitory adipocytes in hips, femurs, and bone marrow, resulting in higher blood counts. Alternatively, WHR-related blood trait variation may reflect inhibitory paracrine or endocrine effects from gluteal or truncal adipose depots (Comazzetto et al., 2021), or differential accumulation of HSC-supportive mesenchymal stem cell-derived bone marrow adipocytes (Zhong et al., 2020).

More broadly, our findings represent the first example of divergent genome-wide effects of generalized obesity (BMI) and truncal adiposity (WHR) on any phenotype. At minimum, this provides a rationale for concurrent BMI and WHR adjustments when analyzing blood trait GWAS loci to avoid directional bias. These adjustments also provide novel stratification criteria for blood trait GWAS fine mapping studies and candidate blood gene selection. This will be particularly important for studied aiming to explain obesity and adiposity-related metabolic or stromal effects on blood cells, which are distinct from cholesterol- or lipid-mediated peripheral effects on erythroid cells at the genetic level.

## Methods

### GWAS summary statistics collection

We analyzed publicly available GWAS summary statistics for blood traits (n=563,085) (Vuckovic et al., 2020), BMI (n=484,680) (Pulit et al., 2019), WHR (n=485,486) (Pulit et al., 2019), and WHRadjBMI (n=484,563) (Pulit et al., 2019), CAD (n=547,261) (Van Der Harst and Verweij, 2018), and lipid traits including TC (n=215,551), TG (n=211,491), LDL (n=215,196), and HDL (n=210,967) (Klarin et al., 2018). Data were derived from individuals of European ancestry only and were analyzed using genome build hg19/GRCh37.

### Instrumental variable creation

To construct instrumental variables (IVs), we identified all SNPs common to all exposure and outcome data sets and clumped genome-wide significant SNPs for the exposure to identify linkage-independent SNPs (EUR r^2^<0.01) in 500 kb regions using TwoSample MR. IV strengths were estimated using F-statistics calculated as described (Burgess and Thompson, 2011). All instrumental variables used in this study can be found on GitHub (https://github.com/thomchr/ObesityAdiposityBloodMR).

### Mendelian randomization and causal effect estimation

Univariable MR analyses (TwoSample MR package v0.5.5 (Hemani et al., 2018)) were conducted using R (v3.6.3). Presented data show causal estimates from inverse variance weighted (IVW, random effects model), weighted median, and MR-Egger regression methods. We assessed pleiotropic bias using MR-Egger regression intercepts, which if significantly non-zero can imply directional bias (Bowden et al., 2015). Multivariable Mendelian randomization analyses utilized the MVMR package (Sanderson et al., 2019) in R. Results shown are IVW method-based causal estimates. Causal direction analyses utilized MR-Steiger and we report values for sensitivity, statistical significance, and inference of the ‘correct causal direction’ (Hemani et al., 2017).

For continuous outcomes (blood traits, lipid traits, BMI), results are presented as beta effect values representing changes in standard deviation units for these traits, per standard deviation unit change in exposure. Standard deviation unit estimations were obtained from Gharahkhani *et al* (Gharahkhani et al., 2019) and Beutler *et al* (Beutler and Waalen, 2006) for BMI and HGB, respectively. For dichotomous outcomes (CAD), causal effect estimates can be converted to odds ratios by exponentiating causal effect estimates (=exp^[effect]) to calculate a value reflecting the change in outcome per standard deviation unit increase in exposure (Burgess and Labrecque, 2018). However, CAD outcome values are presented as SD units to facilitate comparison with blood trait effects.

### Mediation analysis

Mediation analysis estimates were calculated as described (Burgess et al., 2017). Total and direct effects are reported for the exposure and mediating trait on each outcome.

### Conditional GWAS analysis

Conditional analyses of filtered SNP sets, containing SNPs found in BMI, WHR, and blood trait summary statistics, were analyzed using mtCOJO with a limit of r^2^<0.01 (Yang et al., 2011). Results were clumped using plink (v1.90 beta) (Purcell et al., 2007) to identify linkage-independent sentinel SNPs with r^2^<0.01 in 500 kb genomic regions (flagged parameters were -- clump-p1 5E-8 --clump-p2 1 --clump-r2 0.01 --clump-kb 500). Separate experiments were performed on the same filtered SNP sets to adjust for BMI, or both BMI and WHR. To compare uncorrected with BMI- or BMI-and-WHR-adjusted results, we aggregated sentinel SNPs and clumped based on original GWAS p-values (--clump-p1 1 --clump-p2 1 --clump-r2 0.01 --clump- kb 500) to retrieve a complete set of linkage independent loci. This second clump output allowed us to calculate how many regions were shared, nullified, or novel in the adjusted vs unadjusted data sets. The gene nearest to each sentinel locus was identified using bedtools (Quinlan and Hall, 2010). Locus zoom plots were created through the online instrument (http://locuszoom.org, (Pruim et al., 2010)).

### Gene-level analyses

We identified gene and tissue associations for blood trait summary statistics before and after adjustment using FUMA (Watanabe et al., 2019), which uses MAGMA for gene identification (Leeuw et al., 2015).

### Gene Ontology

Gene lists were analyzed for significantly over- or under-enriched GO biological processes. Statistical significance was assigned based on Fisher’s exact test p<0.05 after Bonferroni correction for multiple testing (http://geneontology.org, (Mi et al., 2019)).

### Statistical analyses and data presentation

Estimated effects from exposure(s) on outcome are presented from inverse variance weighted, weighted median, and MR-Egger regression measures. Because Cochran’s Q test (included in the TwoSample MR package (Hemani et al., 2018)) found heterogeneity in some IVs, we utilized the random-effect model when performing inverse-variance weighted MR. Thus, we performed and report MR results using IVs that had not undergone pruning. Statistical significance was defined as P < 0.05 for all experiments. For experiments that analyzed 16 blood traits, we also report those that met a more stringent threshold of p<0.003 (∼0.05/16). Statistics were calculated with GraphPad Prism 8. Figures were prepared using GraphPad Prism 8 and Inkscape (v1.1). Schematic cartoons were created using BioRender.

### Coding scripts and data sets

All relevant coding scripts and data sets can be found on GitHub (https://github.com/thomchr/ObesityAdiposityBloodMR). All data and coding scripts are also available upon request.

## Acknowledgements

CST and BFV designed the study. CST, MW, STC, and BFV conducted, analyzed, and/or interpreted experimental data. CST and BFV wrote the paper. All authors approved the final version of the manuscript.

This work was supported by National Institutes of Health [HD043021 and HL156052 to CST, U01 HL124696 to STC, DK101478 and DK126194 to BFV], a Linda Pechenik Montague Investigator Award (BFV), a Children’s Hospital of Philadelphia K-readiness award (CST), and a NHLBI Progenitor Cell Translational Consortium JumpStart Award (CST).

## Data availability

All coding scripts and data sets can be found on GitHub (https://github.com/thomchr/ObesityAdiposityBloodMR) or by request.

## Figure Supplements

**Figure 1-figure supplement 1.**
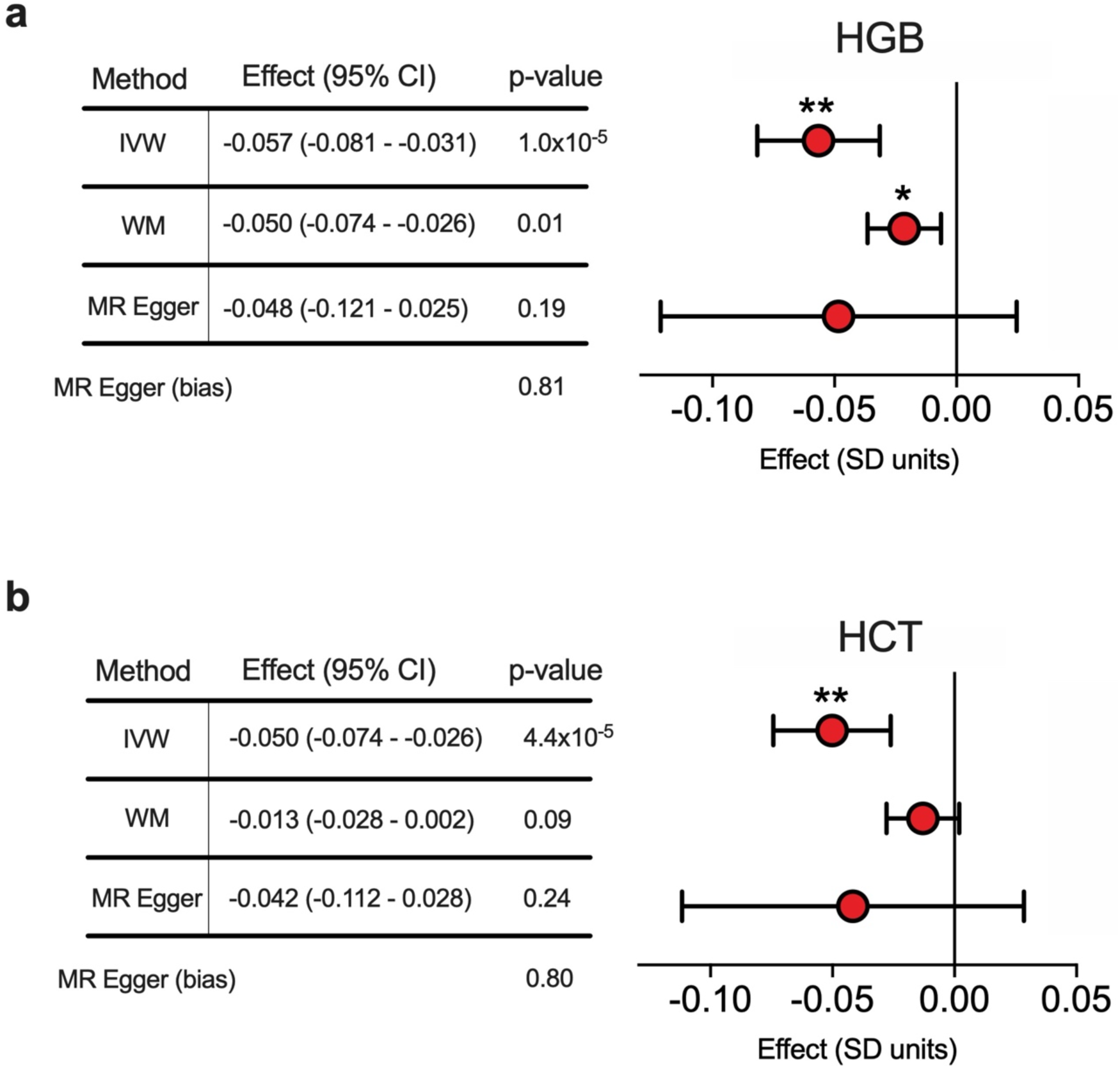
Genetically determined BMI decreases hemoglobin (HGB) and hematocrit (HCT) levels. (**a-b**) Effects of BMI on (**a**) HGB or (**b**) HCT by inverse variance weighted (IVW), weighted median (WM), or MR Egger methods. Effects are in SD units with 95% confidence intervals. Insignificant MR Egger intercept p-values validate effect estimates. *p<0.05.

**Figure 1-figure supplement 2.**
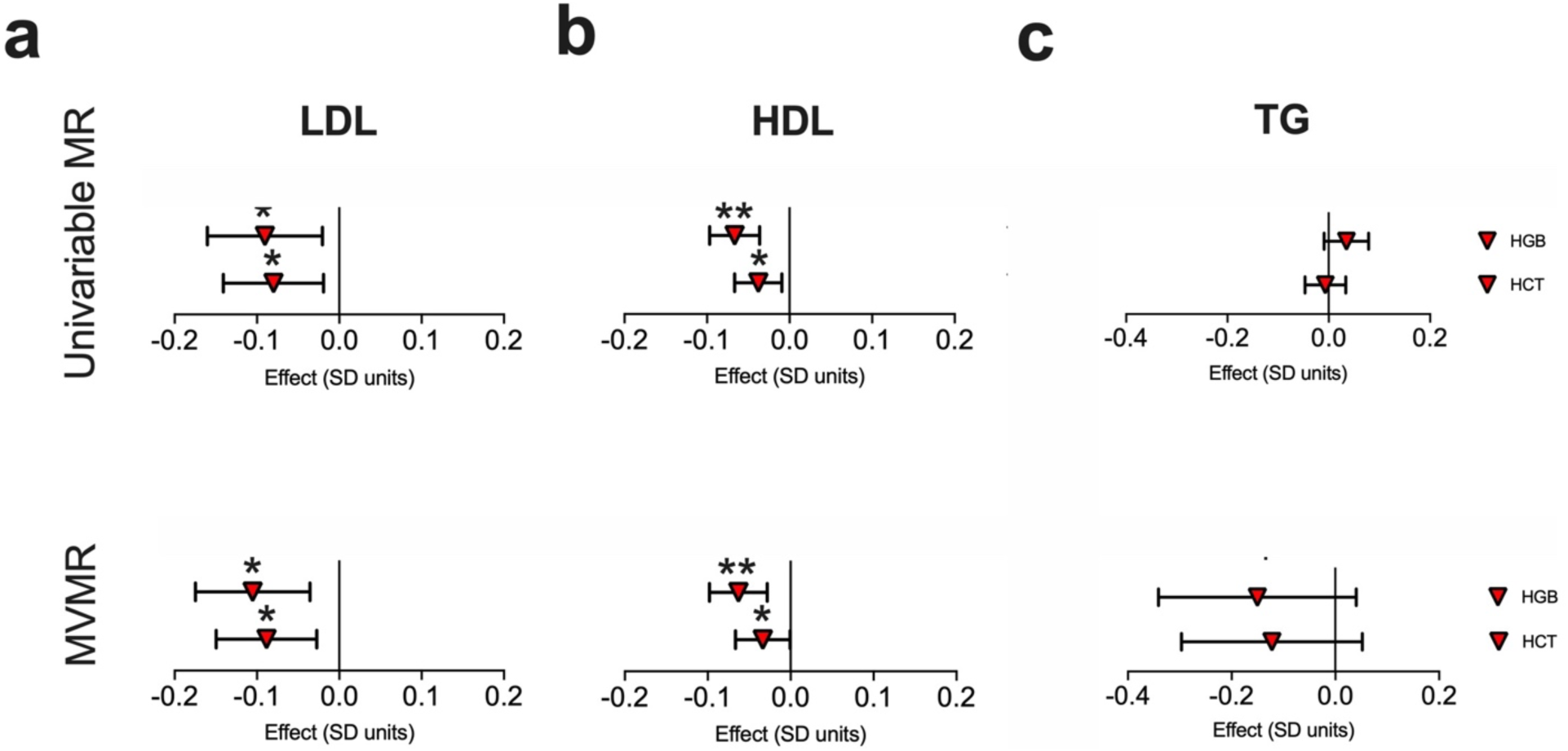
Effects of lipid fractions or triglyceride level on erythroid traits. (**a-c**) Effect sizes (SD units) with 95% confidence intervals for (**a**) low density lipoprotein, LDL, (**b**) high density lipoprotein, HDL, or (**c**) triglyceride level, TG, on erythroid traits (HGB or HCT) by univariable MR (top) or MVMR (bottom). MVMR experiments adjusted for BMI effects. *p<0.05, **p<0.003.

**Figure 1-figure supplement 3.**
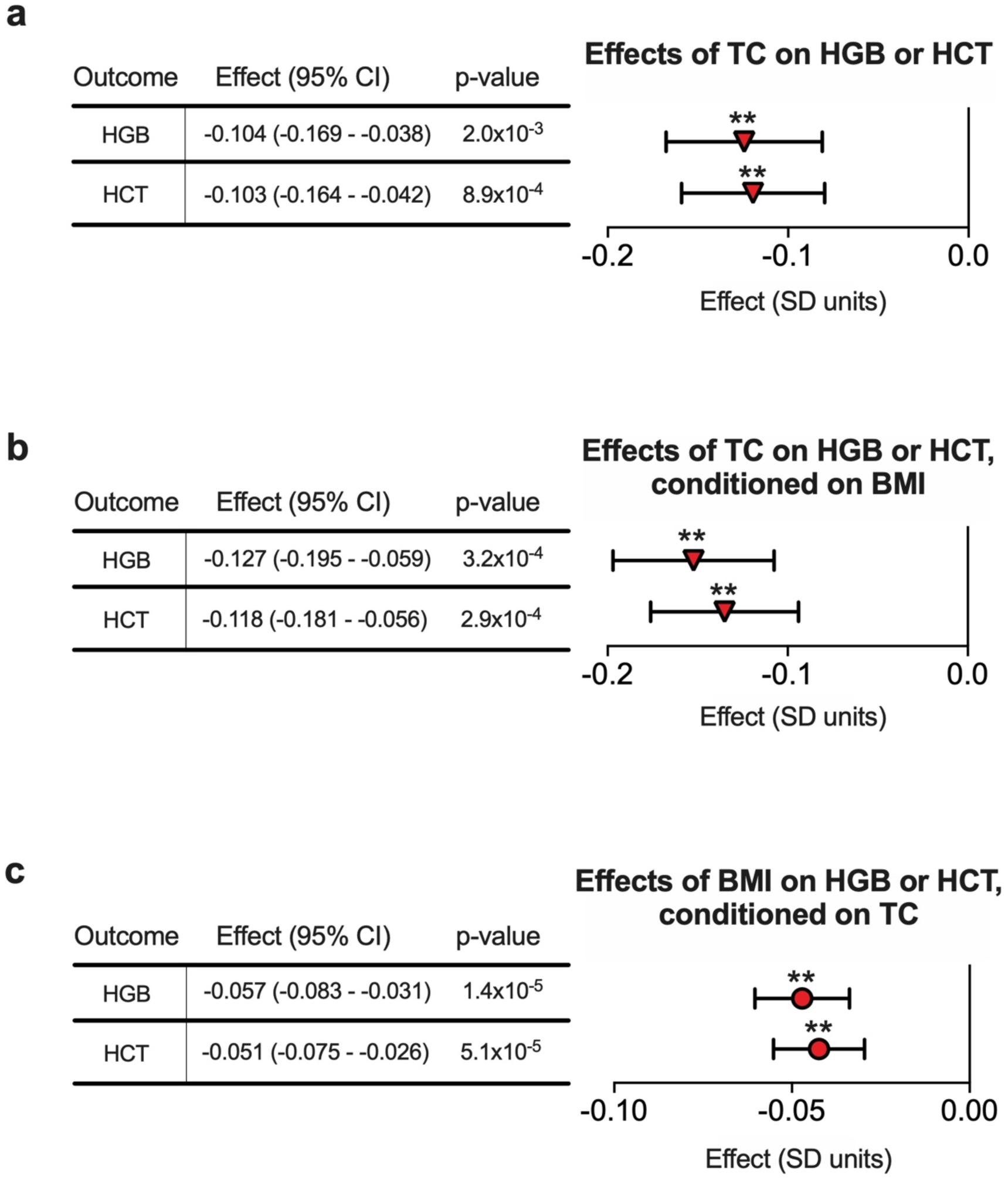
Total cholesterol (TC) decreases HGB and HCT levels independent of BMI effects. (**a**) Effects of TC on HGB or HCT by IVW method. (**b**) Effects of TC on HGB or HCT by IVW after adjusting for BMI. (**c**) Effects of BMI on HGB or HCT by IVW after adjusting for TC. Effects are in SD units, with whiskers showing 95% CI. **p<0.003.

**Figure 1-figure supplement 4.**
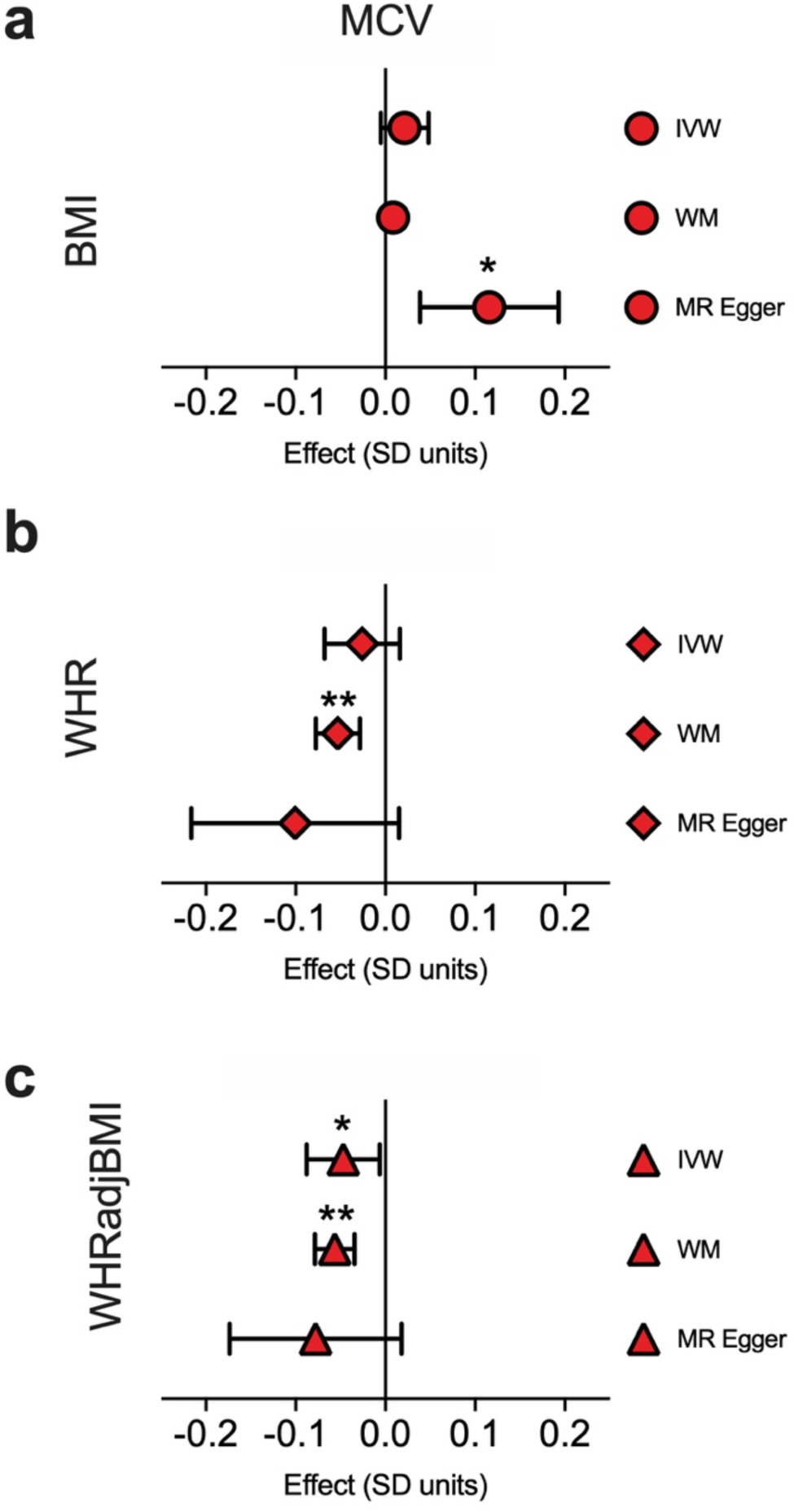
Effects of BMI, WHR, and WHRadjBMI on MCV across MR methodologies. (**a-c**) Effects of (**a**) BMI, (**b**) WHR, or (**c**) WHRadjBMI on MCV by univariable MR. *p<0.05, **p<0.003.

**Figure 1-figure supplement 5.**
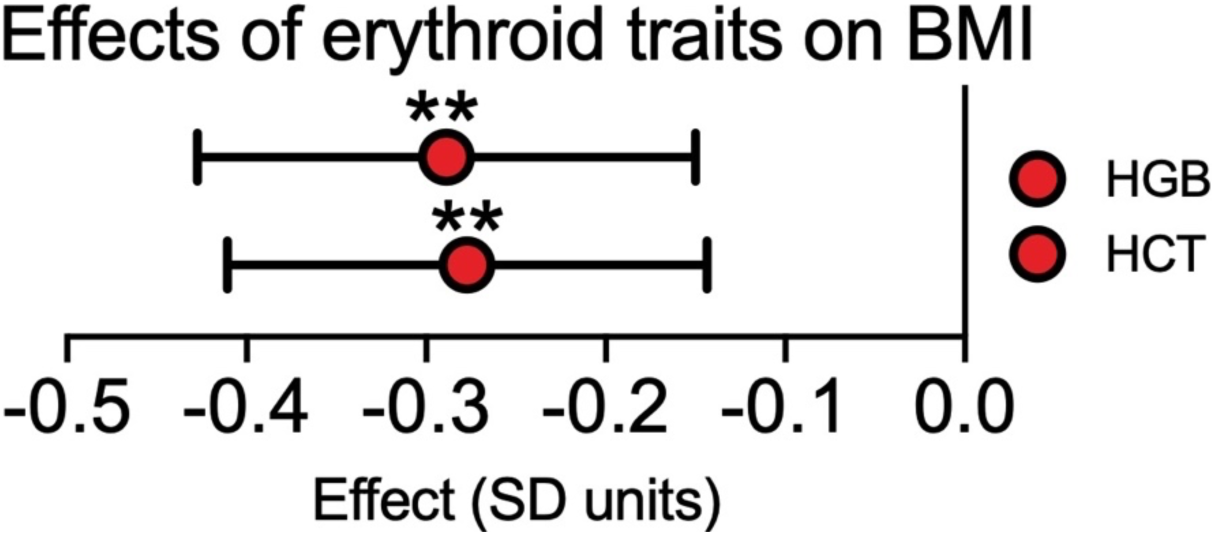
Effects of HGB or HCT on BMI. Two-sample MR experiments show effects (SD units) with 95% confidence intervals for indicated erythroid traits on BMI by IVW method. **p<0.003.

**Figure 1-figure supplement 6.**
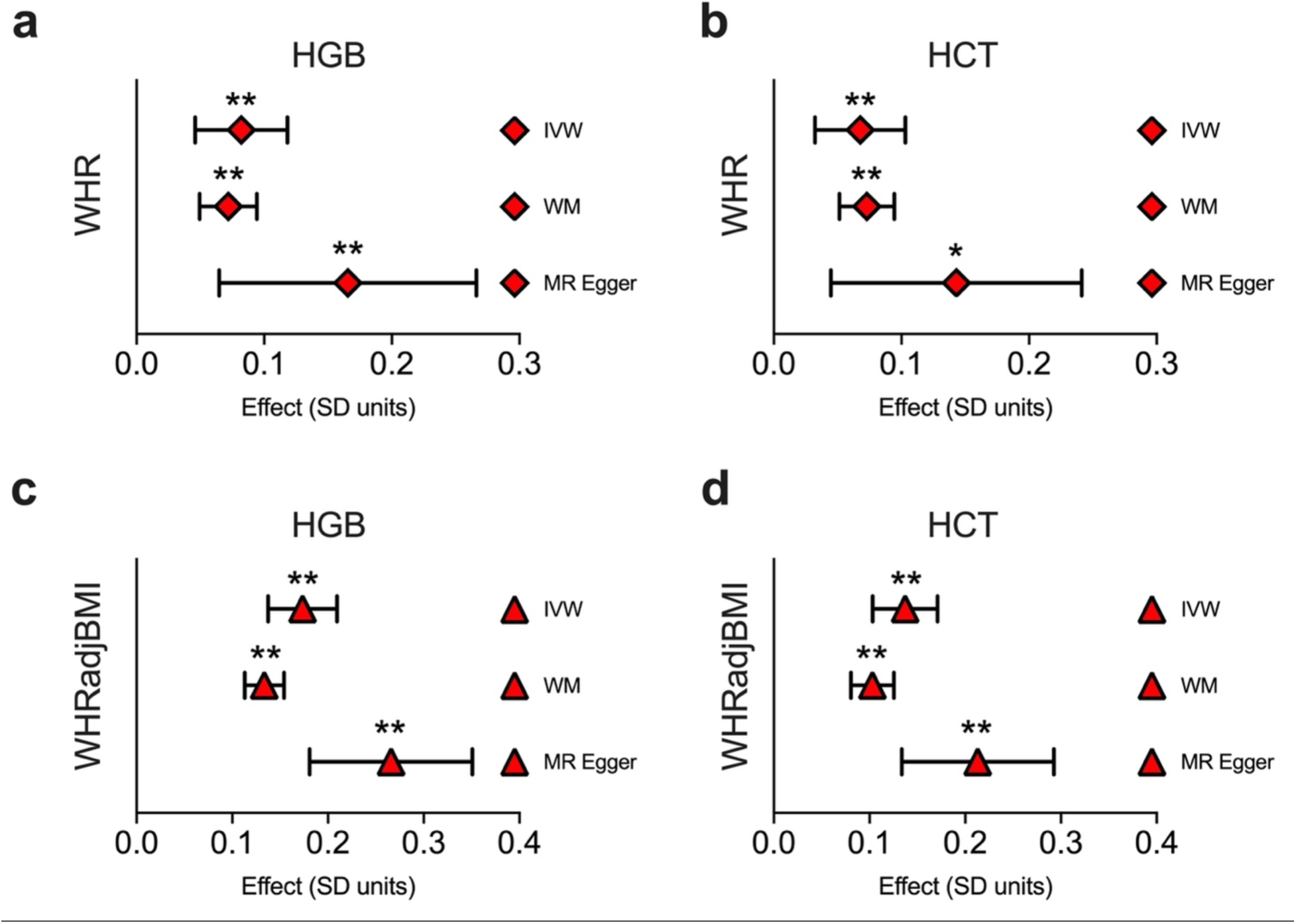
Effects of WHR and WHRadjBMI are consistent across MR methodologies. (**a-b**) By univariable MR, WHR exerts a positive effect on (**a**) HGB and (**b**) HCT. (**c-d**) In analogous experiments, WHRadjBMI exerts a positive effect on (**c**) HGB and (**d**) HCT. Effects were larger after WHR was adjusted for BMI at the individual level (WHRadjBMI). *p<0.05, **p<0.003.

**Figure 1-figure supplement 7.**
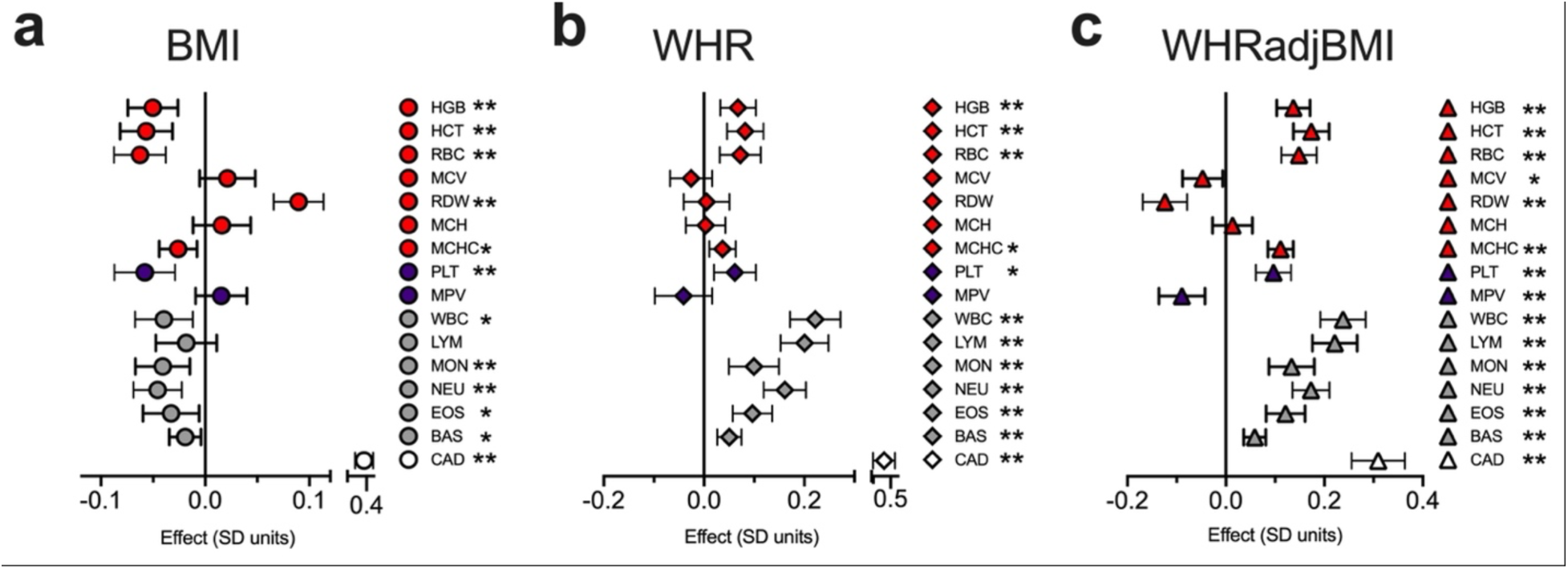
Genetically determined WHR and BMI exert opposing effects on multilineage quantitative blood traits, including red blood cell (RBC), platelet (PLT), and white blood cell (WBC) count. (**a-c**) Univariable MR experiments showing the effects of genetically determined (**a**) BMI, (**b**) WHR, or (**c**) WHRadjBMI on blood traits or coronary artery disease risk (CAD) by inverse variance weighted method (*p<0.05, **p<0.003).

**Figure 1-figure supplement 8.**
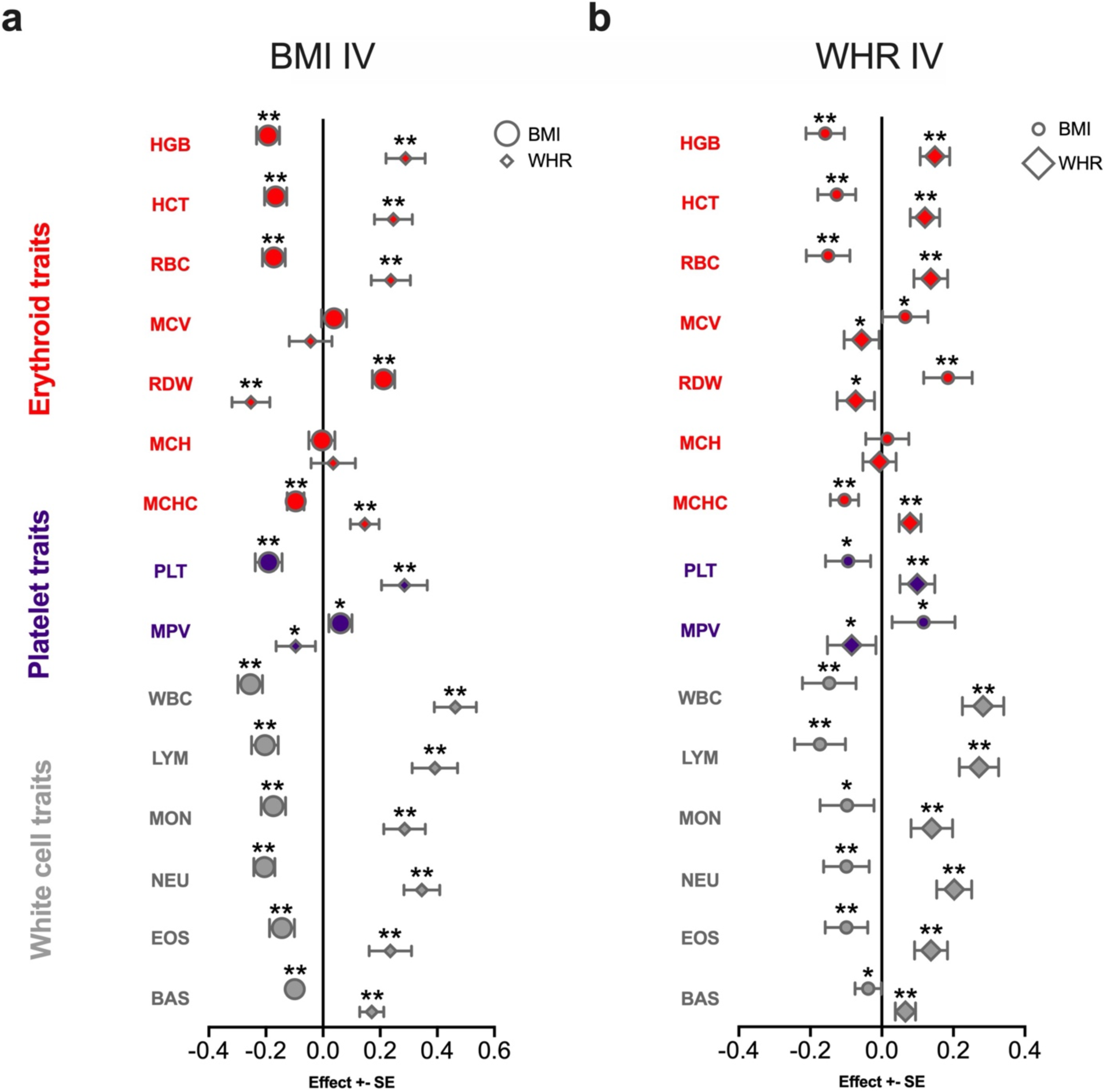
Effects of BMI and WHR on quantitative blood traits by MVMR. (**a-b**) Effects of BMI or WHR, as quantified by MVMR using instrumental variables based on SNPs significant for (**a**) BMI or (**b**) WHR. *p<0.05, **p<0.003.

**Figure 1-figure supplement 9.**
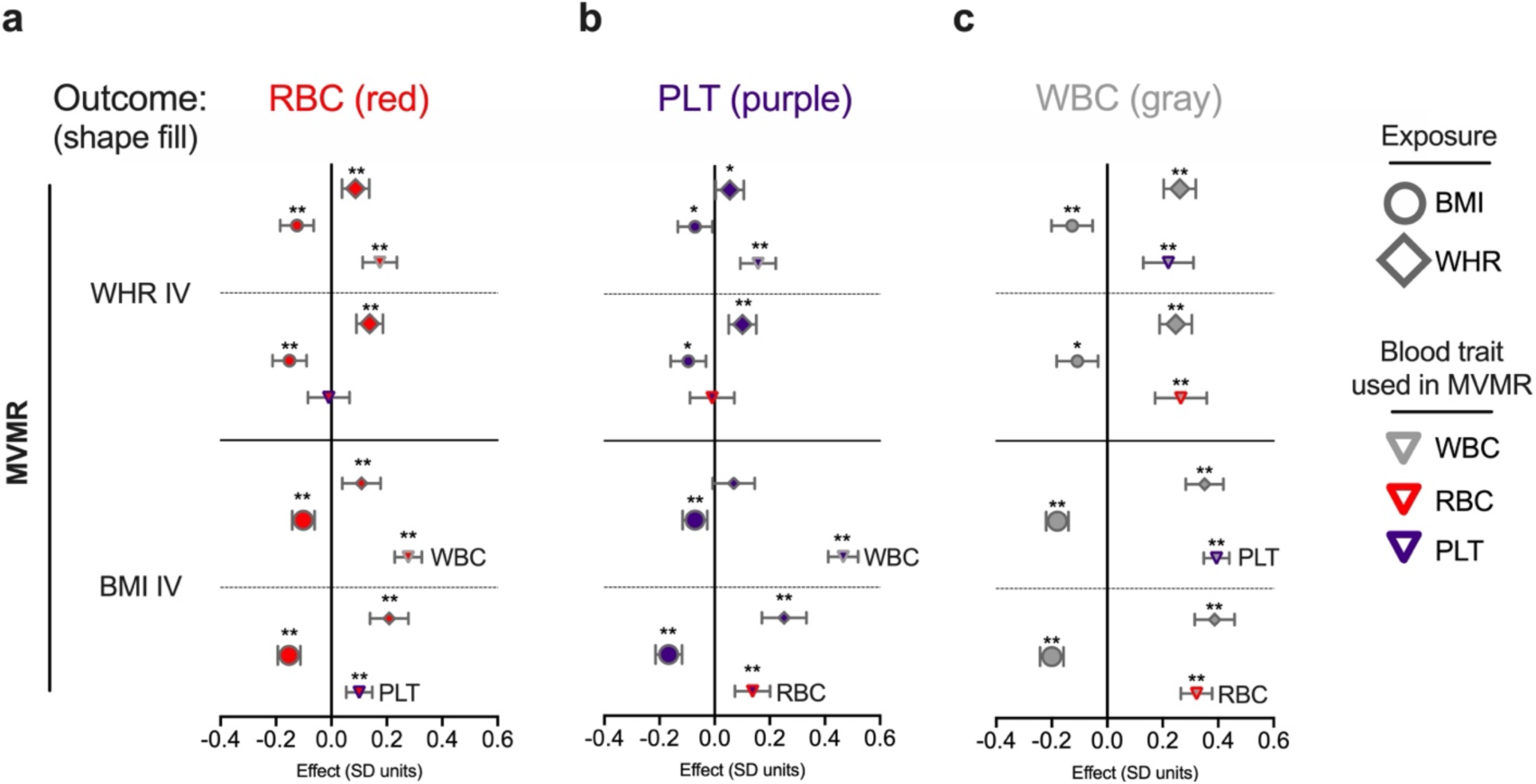
Effects of WHR and BMI on quantitative blood traits by MVMR after regressing out effects of other blood traits. (**a-c**) Effects of WHR, BMI, or WHRadjBMI on (**a**) red blood cell count, RBC, (**b**) platelet count, PLT, or (**c**) white blood cell count, WBC. Left panel indicates trait used to create instrumental variables (639 LD-independent statistically significant SNPs for WHR or 1268 SNPs for BMI). Exposure and mediator traits are listed at right. Within plots, large labels indicate effects from the IV trait with smaller symbols denoting potential mediating traits. *p<0.05, **p<0.003.

**Figure 2-figure supplement 1.**
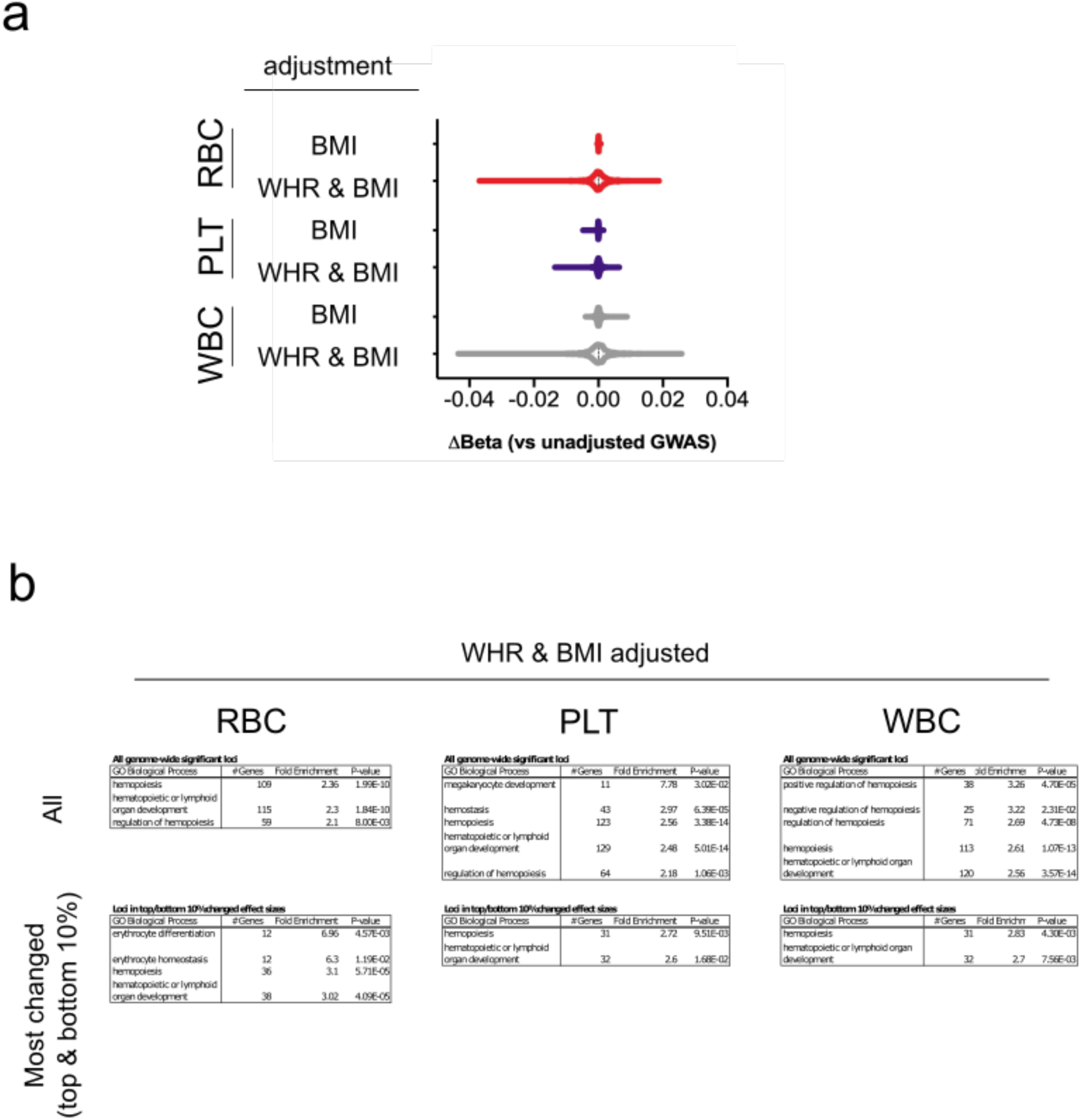
Conditional blood trait analysis based on obesity and/or adiposity modifies interpretation of genomic loci that impact variation in red blood cell (RBC), platelet (PLT), and white blood cell (WBC) counts. (**a**) Violin plots show adjustment for indicated trait(s) via mtCOJO (BMI and/or WHR) modifies effect sizes at genome wide-significant blood trait loci. Adiposity (WHR) disperses effect sizes more than BMI adjustment alone. (**b**) After mtCOJO adjustment for WHR and BMI, significant SNPs and nearby genes excluded blood cell-specific gene ontology pathways. Shown are significant pathways related to hemopoiesis/hematopoiesis, erythrocyte biology, or megakaryocyte/platelet biology (Fisher’s exact test p<0.05 after multiple testing).

**Figure 2-figure supplement 2.**
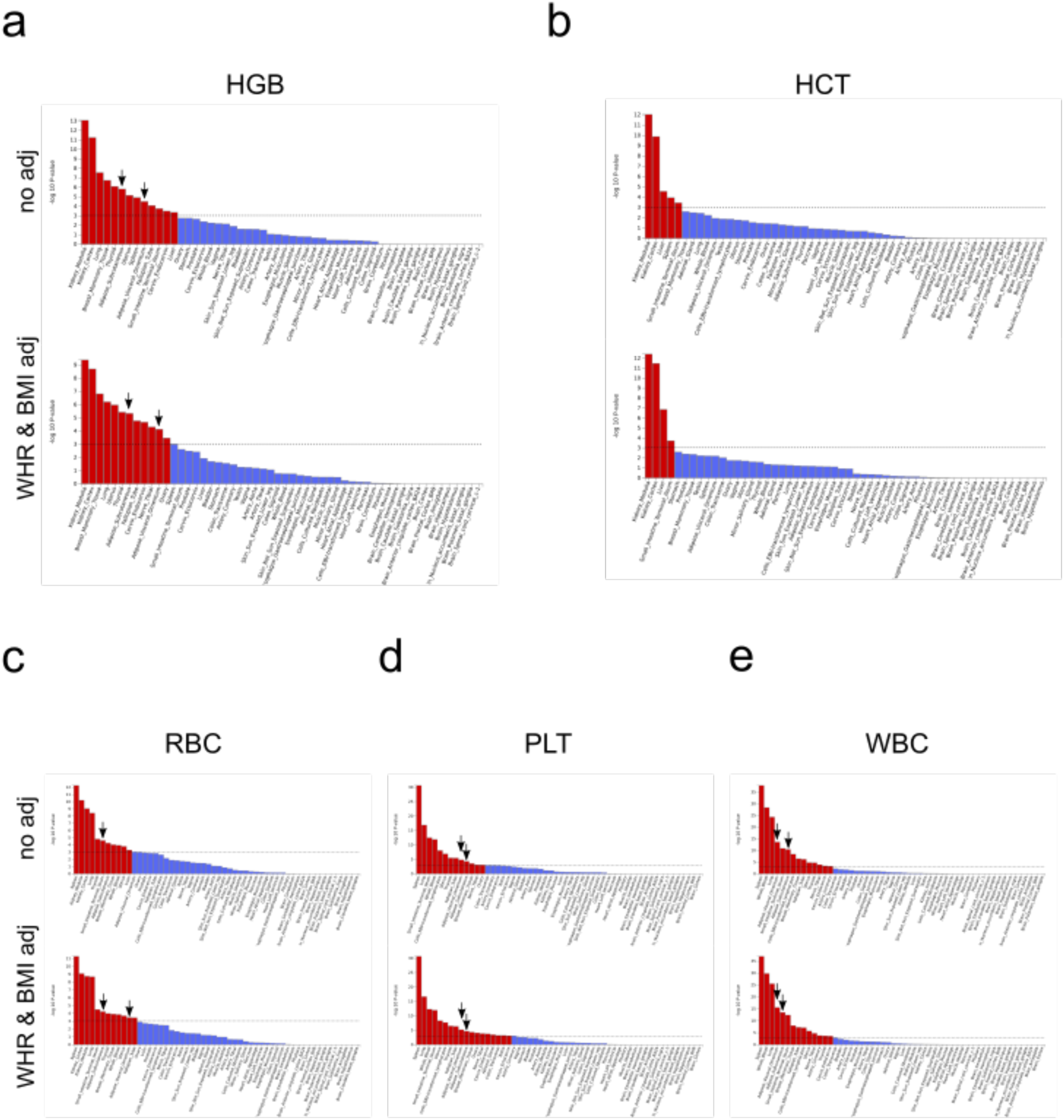
MAGMA tissue enrichment analyses for original and BMI/WHR-adjusted blood trait data. Significantly enriched adipose populations are identified by arrows in each plot.

## Additional file 1

**Table 1.** F statistics and other characteristics for instrumental variables used in this study.

**Table 2.** Mediation analysis results defining total and direct impact of the indicated exposures on HGB or HCT.

**Table 3.** MR Steiger analysis for causal directionality between TC, TG, or BMI versus HGB or HCT.

**Table 4.** New, clarified, shared, or nullified ‘missing’ genome-wide significant loci after adjusting HGB summary statistics for BMI. The nearest gene to each locus is shown.

**Table 5.** New, clarified, shared, or nullified ‘missing’ genome-wide significant loci after adjusting HGB summary statistics for WHR and BMI. The nearest gene to each locus is shown.

**Table 6.** New, clarified, shared, or nullified ‘missing’ genome-wide significant loci after adjusting HCT summary statistics for BMI. The nearest gene to each locus is shown.

**Table 7.** New, clarified, shared, or nullified ‘missing’ genome-wide significant loci after adjusting HCT summary statistics for WHR and BMI. The nearest gene to each locus is shown.

**Tables 8.** Gene ontology analysis of genes nearest to HGB loci before conditional analysis. Presented pathways were significantly enriched by Fisher’s exact test with Bonferroni correction for multiple testing (p<0.05).

**Table 9.** Gene ontology analysis of genes nearest to all genome-wide significant HGB loci after BMI adjustment. Presented pathways were significantly enriched by Fisher’s exact test with Bonferroni correction for multiple testing (p<0.05).

**Table 10.** Gene ontology analysis of genes nearest to select genome-wide significant HGB loci after BMI adjustment. Only loci with effect sizes that were among the top 10% or bottom 10% most affected by BMI adjustment were included in this pathway analysis. Presented pathways were significantly enriched by Fisher’s exact test with Bonferroni correction for multiple testing (p<0.05).

**Table 11.** Gene ontology analysis of genes nearest to all genome-wide significant HGB loci after WHR and BMI adjustment. Presented pathways were significantly enriched by Fisher’s exact test with Bonferroni correction for multiple testing (p<0.05).

**Table 12.** Gene ontology analysis of genes nearest to select genome-wide significant HGB loci after WHR and BMI adjustment. Only loci with effect sizes that were among the top 10% or bottom 10% most affected by BMI adjustment were included in this pathway analysis. Presented pathways were significantly enriched by Fisher’s exact test with Bonferroni correction for multiple testing (p<0.05).

**Tables 13**. Gene ontology analysis of genes nearest to HGB loci before conditional analysis. Presented pathways were significantly enriched by Fisher’s exact test with Bonferroni correction for multiple testing (p<0.05).

**Table 14.** Gene ontology analysis of genes nearest to all genome-wide significant HCT loci after BMI adjustment. Presented pathways were significantly enriched by Fisher’s exact test with Bonferroni correction for multiple testing (p<0.05).

**Table 15.** Gene ontology analysis of genes nearest to select genome-wide significant HCT loci after BMI adjustment. Only loci with effect sizes that were among the top 10% or bottom 10% most affected by BMI adjustment were included in this pathway analysis. Presented pathways were significantly enriched by Fisher’s exact test with Bonferroni correction for multiple testing (p<0.05).

**Table 16.** Gene ontology analysis of genes nearest to all genome-wide significant HCT loci after WHR and BMI adjustment. Presented pathways were significantly enriched by Fisher’s exact test with Bonferroni correction for multiple testing (p<0.05).

**Table 17.** Gene ontology analysis of genes nearest to select genome-wide significant HCT loci after WHR and BMI adjustment. Only loci with effect sizes that were among the top 10% or bottom 10% most affected by BMI adjustment were included in this pathway analysis. Presented pathways were significantly enriched by Fisher’s exact test with Bonferroni correction for multiple testing (p<0.05).

**Table 18.** Gene ontology analysis of genes nearest to all genome-wide significant RBC loci after WHR and BMI adjustment. Presented pathways were significantly enriched by Fisher’s exact test with Bonferroni correction for multiple testing (p<0.05).

**Table 19.** Gene ontology analysis of genes nearest to select genome-wide significant RBC loci after WHR and BMI adjustment. Only loci with effect sizes that were among the top 10% or bottom 10% most affected by BMI adjustment were included in this pathway analysis. Presented pathways were significantly enriched by Fisher’s exact test with Bonferroni correction for multiple testing (p<0.05).

**Table 20.** Gene ontology analysis of genes nearest to all genome-wide significant PLT loci after WHR and BMI adjustment. Presented pathways were significantly enriched by Fisher’s exact test with Bonferroni correction for multiple testing (p<0.05).

**Table 21.** Gene ontology analysis of genes nearest to select genome-wide significant PLT loci after WHR and BMI adjustment. Only loci with effect sizes that were among the top 10% or bottom 10% most affected by BMI adjustment were included in this pathway analysis. Presented pathways were significantly enriched by Fisher’s exact test with Bonferroni correction for multiple testing (p<0.05).

**Table 22.** Gene ontology analysis of genes nearest to all genome-wide significant WBC loci after WHR and BMI adjustment. Presented pathways were significantly enriched by Fisher’s exact test with Bonferroni correction for multiple testing (p<0.05).

**Table 23.** Gene ontology analysis of genes nearest to select genome-wide significant WBC loci after WHR and BMI adjustment. Only loci with effect sizes that were among the top 10% or bottom 10% most affected by BMI adjustment were included in this pathway analysis. Presented pathways were significantly enriched by Fisher’s exact test with Bonferroni correction for multiple testing (p<0.05).

